# Age-dependent neurodegeneration and organelle transport deficiencies in mutant TDP43 patient-derived neurons are independent of TDP43 aggregation

**DOI:** 10.1101/218610

**Authors:** N Kreiter, A Pal, X Lojewski, P Corcia, M Naujock, P Reinhardt, J Sterneckert, S Petri, F Wegner, A Storch, A Hermann

## Abstract

TAR DNA-binding protein 43 (TDP43) is a cause of familiar and sporadic amyotrophic lateral sclerosis (ALS). The diverse postulated mechanisms by which TDP43 mutations cause the disease are not fully understood. Human wildtype and TDP43 S393L and G294V mutant spinal motor neuron cultures were differentiated from patient-derived iPSCs. Mutant hTDP43 and wildtype motor neuron cultures did not differ in neuron differentiation capacity during early maturation stage. During aging we detected a dramatic neurodegeneration including neuron loss and pathological neurofilament abnormalities in TDP43 mutant cultures only. Additionally mitochondria and lysosomes of aging spinal motor neurons revealed robust TDP43 mutation dependent abnormal phenotypes in size, shape, speed and motility which all appeared without TDP43 mislocalization or aggregation formation. Furthermore, D-sorbitol – known to induce stress granules and cytoplasmic mislocalization of TDP43 – rescued axonal trafficking phenotypes without any signs of TDP43 mislocalization or aggregation formation. Our data indicate TDP43 mutation-dependent but cytosolic aggregation-independent mechanisms of motor neuron degeneration in TDP43 ALS.

## Introduction

Worldwide 2-4 of 100.000 people per year suffer from amyotrophic lateral sclerosis (ALS) which is the most frequent variant of motor neuron disease with a survival time of 1-5 years after symptom onset. Motor neurons are the mainly affected cell type that undergoes degeneration and death in ALS ^1,2^. Several gene mutations were found to cause familiar forms of ALS (fALS) of which the first identified were *SOD1* (superoxide dismutase 1) gene mutations in 1993 ^3,4^. In 2006 mutations in *TDP43* (TAR DNA binding protein, TDP43) were identified to cause fALS as well as sporadic ALS but the pathophysiology caused by mutant TDP43 is still not fully understood ^5^. Most importantly, TDP43 is the main aggregating protein in sporadic ALS also being aggregated in some fALS forms (e.g. TDP43, C9ORF72) ^6^. However, TDP43-mediated pathophysiology appears to be mechanistically distinct from mutant SOD1 action ^7^.

Different mechanisms for TDP43 mutant pathology have been suggested (for review see Jovicic). These include nuclear loss of function leading to transcription and splicing defects, toxic properties in the cytoplasm sequestering RNAs and RNA binding proteins and finally loss of axonal transport function with RNA granule transport deficiencies ^8^. The main focus has been drawn on pathological accumulation of cytosolic TDP43 ^9-11^. Mutant TDP43 human induced pluripotent stem cell (hiPSC)-derived neurons were reported to show elevated levels of soluble and detergent-resistant TDP43 protein and decreased survival in long-term differentiation ^12^. Such lines showed a two-fold increase in cytosolic TDP43 compared to the controls which could be – at least partially – reduced by 30% by allele-specific knockdown ^13^. However, on a closer look, also in models reporting cytosolic mislocalization of TDP43, only up to one third of cells showed this phenotype while the remaining cells still showed physiological nuclear TDP43 ^10,11,14^.

TDP43 was shown to be actively transported within the axon in both directions ^14^. TDP43 thereby forms cytoplasmic mRNP granules that undergo bidirectional, microtubule-dependent transport in neurons and facilitate delivery of target mRNA to distal neuronal compartments ^15^. TDP43 mutations interfere with mRNA transport function ^14,15^. Neurofilament light chain (NEFL) mRNA was shown to be transported in such TDP43 granules and was reduced in TDP43 mutant motor neurons ^10,15^.

ALS-linked mutations in TDP43 can impair stress granule dynamics ^16^. TDP43 had previously been shown to associate with multiple proteins that are part of mRNP granules (e.g., staufen, FMRP, SMN, and HuD), including neuronal transport granules ^17^. TDP43 was shown to be directed to stress granules by sorbitol, a novel physiological osmotic and oxidative stressor ^18^ and to regulate FOXO-dependent protein quality control in stress response ^19^.

Mutations in TDP43 led to abnormal ^9^ or reduced neurite outgrowth ^20,21^. TDP43 mice model shows lack of mitochondria in motor axon terminals ^10^. Furthermore, TDP43 was reported to impair mitochondria and lysosome morphology and function in cell models of ALS. Interestingly, both overexpression and knockdown caused perturbations in primary murine neurons suggesting a tight regulation of TDP43 ^22,23^. Overexpression of wildtype or mutant TDP43 caused mitochondrial shortening in dendrites, not in axons, due to increased fission and decreased mitochondrial movement in axons and dendrites. In contrast, knockdown increased mitochondrial length but also decreased dendritic and axonal mitochondrial motility. Mt TDP43 caused co-staining of mitochondria and TDP43, which was rescued by Mfn2 expression.

However, the relation between (stress induced) cytoplasmic TDP43 aggregation and neuronal dysfunction and degeneration remains enigmatic. Furthermore, data on organelle trafficking and neurite morphology in human patient-derived motor neurons are still lacking.

This prompted us to use patient-derived induced pluripotent stem cells (hiPSCs)-derived motor neurons to investigate (i) the sequential appearance of neuronal dysfunction and neurodegeneration, (ii) aggregate formation and (iii) how these events are mechanistically connected. Surprisingly, we found significant neuronal dysfunction and degeneration during cellular aging, but did not find relevant signs of pathological TDP43 aggregation, arguing against an upstream function of TDP43 aggregation in TDP43-ALS.

## Materials and Methods

### Patient characteristics

We included cell lines carrying a “benign” (S393L, late onset primary anarthria with ALS/LMND, no clinical symptoms of FTD, female, family history of ALS and PD) and a “malign” (G294V, early onset ALS, no clinical symptoms of FTD, male, no family history) TDP43 mutation and were compared to four wildtype cell lines from healthy volunteers (female, age at biopsy 45; female, age at biopsy 53; male, age at biopsy 60 ^24,25^). An overview of the used cell lines is demonstrated in Supplemental Table S1. The performed procedures were in accordance with the Helsinki convention and approved by the Ethical Committee of the University of Dresden (EK45022009; EK393122012). Informed consent was obtained from any individual including informed consent about publishing data obtained from the iPSCs derived from skin fibroblasts. The skin biopsies (see below) were obtained after anonymisation.

### Generation of iPSC lines

For generating iPSC lines fibroblast lines were established from skin biopsies taken from familial ALS patients and healthy controls. The generation and characterization of wildtype iPSC lines was reported previously ^24,26–28^. Briefly, patient fibroblasts were reprogrammed using pMX-based retroviral vectors encoding the human cDNAs of OCT4, SOX2, KLF4 and cMYC (pMX vectors). Vectors were co-transfected with packaging-defective helper plasmids into 293T cells using Fugene 6 transfection reagent (Roche Diagnostics). Fibroblasts were plated at a density of 50,000 cells/well on 0.1% gelatin-coated 6-well plates and infected three times with a viral cocktail containing vectors expressing OCT4:SOX2:KLF4:cMYC in a 2:1:1:1 ratio in presence of 6 µg/ml protamine sulfate (Sigma Aldrich) and 5 ng/ml FGF2 (Peprotech). Infected fibroblasts were plated onto mitomycin C (MMC, Tocris) inactivated CF-1 mouse embryonic fibroblasts (in-lab preparation) at a density of 900 cells/cm2 in fibroblast media. The next day media was exchanged to ES medium containing 78% Knock-out DMEM, 20% Knock-out serum replacement, 1% non-essential amino acids, 1% penicillin/streptomycin/glutamine and 50µM β-Mercaptoethanol (all from Invitrogen) supplemented with 5 ng/ml FGF2 and 1 mM valporic acid (Sigma Aldrich). Media was changed every day to the same conditions. iPSC-like clusters started to appear at day 7 post infection, were manually picked 14 days post-infection and plated onto CF-1 feeder cells in regular ES-Media containing 5 ng/ml FGF2. Stable clones were routinely passaged onto MMC-treated CF-1 feeder cells (Globalstem) using 1 mg/ml collagenase type IV (Invitrogen) and addition of 10 µM Y-27632 (Ascent Scientific) for the first 48 hours after passaging. Media change with addition of fresh FGF2 was performed every day.

Stable clones were analyzed by qRT-PCR for silencing of viral transgenes prior to further experimental procedures.

### Trilineage differentiation potential

iPSC colonies were grown under standard conditions, cleaned and treated with collagenase type IV (2mg/ml, Invitrogen). Floating aggregates were collected and transferred into ultra-low attachment plates (NUNC) in regular ES-Media containing 5 µM Y-27632 (Ascent Scientific) for meso-/endodermal differentiation or ES-Media containing 5 µM Y-27632, 10 µM SB431542 (Tocris) and 1 µM Dorsomorphin (Tocris) for ectodermal differentiation. Two days later the medium was changed to the same conditions leaving out the Y-27632. After four days of EB formation, aggregates were plated onto gelatin (0.1%, Millipore) coated wells for meso-/endodermal differentiation or onto plates coated with MatrigelTM (BD Bioscience) for ectodermal differentiation. EBs were differentiated for two weeks using 77.9% DMEM (high glucose, Invitrogen), 20% FCS (PAA), 1% non-essential amino acids (Invitrogen), 1% penicillin/streptomycin/glutamine (Invitrogen) and 0.1% β-Mercaptoethanol (Invitrogen) for the meso-/endodermal lineage and 50% DMEM/F12 (Invitrogen), 50% Neurobasal (Invitrogen) containing 1:200 N2 supplement (Invitrogen), 1:100 B27 supplement without vitamin A (Invitrogen), 1% penicillin/streptomycin/glutamine, 0.1% β-Mercaptoethanol and 1:500 BSA Fraction V (Invitrogen) for ectodermal differentiation.

### Karyotyping

TDP43 iPSC and wildtype cell lines were karyotyped using the HumanCytoSNP-12v array. All clones showing pathological SNPs were excluded (data not shown).

### Genotyping

TDP43 iPSC lines were genotyped after all other characterization had been finished. This was done by a diagnostic human genetic laboratory (CEGAT, Tübingen, Germany) using diagnostic standards.

### Differentiation of human NPCs to spinal motor neurons

The generation of human NPCs and motor neurons was accomplished following the protocol from Reinhardt et al. ^26^. In brief colonies of iPSCs were collected and stem cell medium was added containing 10µM SB-431542, 1µM Dorsomorphin, 3µM CHIR 99021 and 0.5µM purmorphamine ^29^. After 2 days hESC medium was replaced with N2B27 consisting of the aforementioned factors and DMEM-F12/ Neurobasal 50:50 with 1:200 N2 Supplement, 1:100 B27 lacking Vitamin A and 1% penicillin/ streptomycin/ glutamine. On day 4 150µM Ascorbic Acid was added while Dorsomorphin and SB-431542 were withdrawn. 2 Days later the EBs were mechanically separated and replated on Matrigel coated dishes. For this purpose Matrigel was diluted 1:100 in DMEM-F12 and kept on the dishes over night at room temperature. Possessing a ventralized and caudalized character the arising so called small molecule NPCs (smNPC) formed homogenous colonies during the course of further cultivation. It was necessary to split them at a ratio of 1:10 – 1:20 once a week using Accutase for 10 min at 37°C.

Final motor neuron differentiation was induced by treatment with 1µM PMA in N2B27 exclusively. After 2 days 1µM retinoic acid (RA) was added. On day 9 another split step was performed to seed them on a desired cell culture system. Furthermore the medium was modified to induce neuronal maturation. For this purpose the developing neurons were treated with N2B27 containing 10 ng/ µl BDNF, 500 µM dbcAMP and 10 ng/ µl GDNF. Following this protocol it was possible to keep the cells in culture for over 2 months. The cells were analyzed at day 5 and day 32 post maturation induction by immunofluorescence imaging.

### Immunofluorescence of spinal motor neurons

For immunofluorescence staining, cells were washed twice with PBS without Ca2^+^/Mg2^+^ (LifeTechnologies) and fixed with 4% PFA in PBS for 15 min at RT. PFA was aspirated and cells were washed three times with PBS. Fixed cells were first permeabilized for 10 minutes in 0.2 % Triton X solution and subsequently incubated for 1 hour at RT in blocking solution (5% donkey serum in PBS). Following blocking, primary antibodies were diluted in PBS and cells were incubated with primary antibody solution overnight at 4°C. The following primary antibodies were used: chicken anti-SMI32 (1:10000, Covance), rabbit anti-beta-III-Tubulin (1:3000, Covance), rabbit anti-CHAT (1:1500, Chemicon), mouse anti-MAP2 (1:500, BD Biosciences). Post to the primary antibodies cells were washed three times for 5 min with PBS. Secondary antibodies were diluted in PBS and incubated with the cells for 1.5 hours at RT. Following secondary antibodies were used: donkey anti-chicken IgY FITC (1:500, Merck Millipore), donkey anti-rabbit IgG647 (1:500, Life Technologies), donkey anti-rabbit IgG488 (1:500, Life Technologies) and donkey anti-mouse IgG555 (1:500, Life Technologies). Nuclei were counter stained using Hoechst (LifeTechnologies).

### Microfluidic chambers

The MFCs were purchased from Xona (RD900). At first, Nunc glass bottom dishes with an inner diameter of 27 mm were coated with Poly-L-Ornithin over night at 37°C. Following 3 washing steps with sterile water they were kept under the sterile hood to dry out completely. MFCs were sterilized with 70% Ethanol and also left drying. Next the MFCs were dropped on the glass bottom dishes and carefully pressed on the surface. Properly attached the system was then perfused with 0.01 mg/ ml Laminin in PBS. After another 1h at 37° and one washing step with medium it could be used for cell culture. For seeding cells into this system, the entire volume was soaked out and 10µl containing a high concentration of cells (15×10^6 cells/ ml) were directly injected into the main channel connecting two wells. After allowing attachment for half an hour in the incubator, the still empty wells were filled up with maturation medium. This method grants the possibility to enhance the amount of neurons growing in main channels while the wells stay free of them which reduces the medium turnover to a minimum. To avoid drying out PBS was added around the MFCs. Two days later the medium was replaced in a manner which should give the neurons a guidance cue to grow through the microchannels. Therefore a gradient style was established which was accomplished by adding 100 µl N2B27 with 500µM dbcAMP to the seeding site and 200µl N2B27 with 500µM dbcAMP, 10 ng/ µl BNDF, 10 ng/ µl GDNF and 100 ng/ µl NGF to the exit site. The medium composition was replaced every third day. After 7 days the first axons began spreading out at the exit site. It was possible to keep them in this system for up to 6 weeks. MFCs were analyzed by live cell imaging at 14 and 32 days post maturation induction. At day 32 they were treated with 40 mM D-sorbitol in medium for 24h following live cell imaging.

### Live cell imaging

For tracking of lysosomes and mitochondria, cells were double-stained live with 50nM Lysotracker Red DND-99 (Molecular Probes Cat. No. L-7528) and 50nM Mitotracker Deep Red FM (Molecular Probes Cat. No. M22426). For measuring the mitochondrial membrane potential (and tracking as well), cells were stained with 200nM Mitotracker JC-1 (Molecular Probes Cat. No. M34152). Trackers were directly added to culture supernatants and incubated for 1 hr at 37°C. Imaging was then performed without further washing of cells. Live imaging of compartmentalized axons in Xona Microfluidic Chambers (MFC) was performed with a Leica HC PL APO 100x 1.46 oil immersion objective on an inversed fluorescent Leica DMI6000 microscope enclosed in an incubator chamber (37°C, 5% CO2, humid air) and fitted with a 12-bit Andor iXON 897 EMCCD camera (512*512, 16µm pixels, 229.55 nm/pixel at 100x magnification). For more details, refer to https://www.biodip.de/wiki/Bioz06_-_Leica_AFLX6000_TIRF. Excitation was performed with a TIRF Laser module in epifluorescence (widefield) mode with lines at 561nm and 633nm. Fast dual color movies were recorded at 3.3 frames per second (fps) per channel over 2 min (400 frames in total per channel) with 115 ms exposure time as follows: Lysotracker Red (excitation: 561nm, emission filter TRITC 605/65 nm) and Mitotracker Deep Red (excitation: 633nm, emission filter Cy5 720/60 nm). Movie acquisition was performed at strictly standardized readout positions within the micro channels of the micro groove barrier that separated the proximal seeding site from the distal axonal exit. Specifically, the readout windows were located either just adjacent to the channel exit (distal readout) or the channel entry (proximal readout).

### Tracking analysis

Movies were analyzed with FIJI software using the TrackMate v2.7.4 plugin for object (lysosomes and mitochondria) recognition and tracking. Settings were as follows: Pixel width: 0.23µm, Pixel height: 0.23µm, Voxel Depth: 1µm, Crop settings: not applied, Select a detector: DoG detector with estimated blob size: 1.6µm, Threshold: 45, median filter: no, subpixel localization: yes, Initial thresholding: none, Select view: HyperStack Displayer, Set filters on spots: quality above 45, Select a tracker: Linear Motion LAP tracker, Initial search radius. 2µm, Search radius: 2µm, Max. frame gap: 2, Set filters on tracks: track duration ≥ 3 sec. Typically, 200-500 tracks per movie were obtained and analyzed with respect to track displacement (measure for processive, i.e. straight, motility as opposed to undirected random walks) and mean speed. Results were assembled and post-filtered (threshold for track displacement ≥ 1.2µm) in KNIME and MS Excel and bulk statistics analyzed and displayed as box plots in GraphPad Prism 5 software. A minimum of 5 movies (showing 2 micro channels each) was acquired at each readout positions (distal versus proximal) per line, condition and experiment resulting in a minimum of 15 movies in total for the batch analysis.

#### Static analysis of cell organelles

For analysis of organelle count and morphology (mitochondria: elongation; lysosomes: diameter), object segmentation, thresholding and shape analysis was performed with a sequence of commands in FIJI software executed with Macro1 for mitochondria:

run("Clear Results"); run("Z Project…", "start=1 stop=400 projection=[Min Intensity]"); run("Enhance Contrast…", "saturated=0.1 normalize"); run("Tubeness", "sigma=0.4 use"); run("Enhance Contrast…", "saturated=0.1 normalize"); run("8-bit"); setAutoThreshold("Triangle dark"); run("Convert to Mask"); run("Skeletonize (2D/3D)"); run("Set Measurements…", "area redirect=None decimal=5"); run("Create Selection"); run("Measure"); saveAs("Measurements", path+name+" network.txt"); selectWindow(name); run("Slice Keeper", "first=1 last=1 increment=1"); run("Select All"); run("Set Measurements…", "kurtosis redirect=None decimal=5"); run("Clear Results"); run("Measure"); saveAs("Measurements", path+name+" Kurtosis.txt"); run("Grays"); run("Enhance Contrast…", "saturated=0.1 normalize"); run("Subtract Background…", "rolling=3"); setAutoThreshold("Moments dark"); run("Threshold…"); run("Convert to Mask"); run("Set Measurements…", "area fit shape feret's integrated area_fraction limit redirect=None decimal=5"); run("Clear Results"); run("Analyze Particles…", "size=4-Infinity pixel circularity=0.00-1.00 show=Ellipses clear display")

and Macro2 for lysosomes:

run("Clear Results"); run("Slice Keeper", "first=1 last=1 increment=1"); run("Grays"); run("Enhance Contrast…", "saturated=0.1 normalize"); run("Subtract Background…", "rolling=5"); setAutoThreshold("Moments dark"); run("Threshold…"); run("Convert to Mask"); run("Set Measurements…", "area fit shape feret’s integrated area_fraction limit redirect=None decimal=5"); run("Analyze Particles…", "size=3-Infinity pixel circularity=0.00-1.00 show=Ellipses clear display")

These macros returned result tables containing the aspect ratio of fitted eclipses (long:short radius) that was taken as measure for mitochondrial elongation as well as the outer Feret’s diameter that was taken as lysosomal diameter. The same set of movies as for the tracking analysis (see above) was used (first frame only). Typically, hundreds of organelles were analyzed per movie. For bulk statistics, the same batch analysis as for the tracking analysis was performed with resultant distributions displayed as box plots.

### Electrophysiology

We performed patch-clamp recordings as described previously ^30^. All electrophysiological experiments were recorded during week 7 of total differentiation. In brief, we plated 300k cells per Matrigel-coated coverslip in a 24 well plate on day 25. To make sure we recorded from MNs we selected large (>20pF) neurons with multiple neurites only. We also filled the internal patch solution with secondary antibody Alexa 488 to allow MN identification after an additional immunostaining step which gave us 90.5% positively identified MNs ^25^.

### Quantification and statistics

Randomly assigned images of different experiments were quantified on day 14 of neuronal differentiation (=day 5 of neuron maturation) to evaluate MN differentiation capacity. Statistical evaluation was performed using 1-way ANOVA with Bonferroni *post hoc* test for multiple comparison. Data are depicted as mean ± standard deviation from at least four independent experiments each.

Static and dynamic live cell imaging box plot statistics are provided in Suppl. Tables 2-9. Box plot settings: boxes from 25-75 percentile, whiskers from 1-99 percentile, outliers as dots, median as horizontal center line, mean as cross. Differences between conditions (i.e. cells lines, compound treatments, etc.) were revealed with the non-parametric Kruskal-Wallis test including Dunn’s *post hoc* test for non-Gaussian distributions with a significance level of P≤0.05 and 95% confidence interval. It was used to compute significant mean rank differences for the distribution of measured organelle property data in order to evaluate the behavior of the bulk majority of organelles, respectively. Box plots represent batch results merged from all mutant ALS-TDP43 lines, respectively, from at least three independent experiments. Detailed box plot statistics are presented in Supplemental Tables 2-9.

## Results

### Motor neuron differentiation is not affected by ALS-causing TDP43 mutations

In order to confirm the neuronal differentiation potential and ensure that wildtype and TDP43 mutant motor neuron cultures exhibit the same differentiation capacity for further disease modeling, they were analyzed towards the proportion of neuron types, their morphology and their survival rates during motoneuronal differentiation. For this, iPSCs were derived from two different patients and three different control individuals (different families with different mutations; G294V; S393L, respectively; different controls from different families, see also Suppl. Table 1). A defined cell number of patient NPCs (neural progenitor cells) were differentiated for 9 days and subsequently seeded for the final motor neuron maturation process for 5 additional days (Figure 1 A, schematic differentiation procedure). There was no difference in neuron morphologies between wildtype and TDP43 mutant neurons. The calculation of neuron types resulted in equal numbers of either βIII-Tubulin (pan-neuronal marker, young neurons), MAP2 (mature neurons) or SMI32 (motor neurons) positive neurons when wildtype and TDP43 mutants were compared (Figure 1 B-E, *P*=0.107, *F*=2.428). Nearly all neurons marked by βIII-Tubulin were also positive for the mature neuron marker MAP2 (wildtypes: 96.90±1.06 %; TDP43 S393L: 98.5±3.0 %; TDP43 G294V: 100.5±1.0). The majority of mature MAP2^+^ neurons were also identified as SMI32^+^ spinal motor neurons in wildtypes and TDP43 mutants (wildtypes: 60.4±5.3 %; S393L: 68.3±4.1 %; G294V: 71.1±3.5 %).

**Figure 1:**
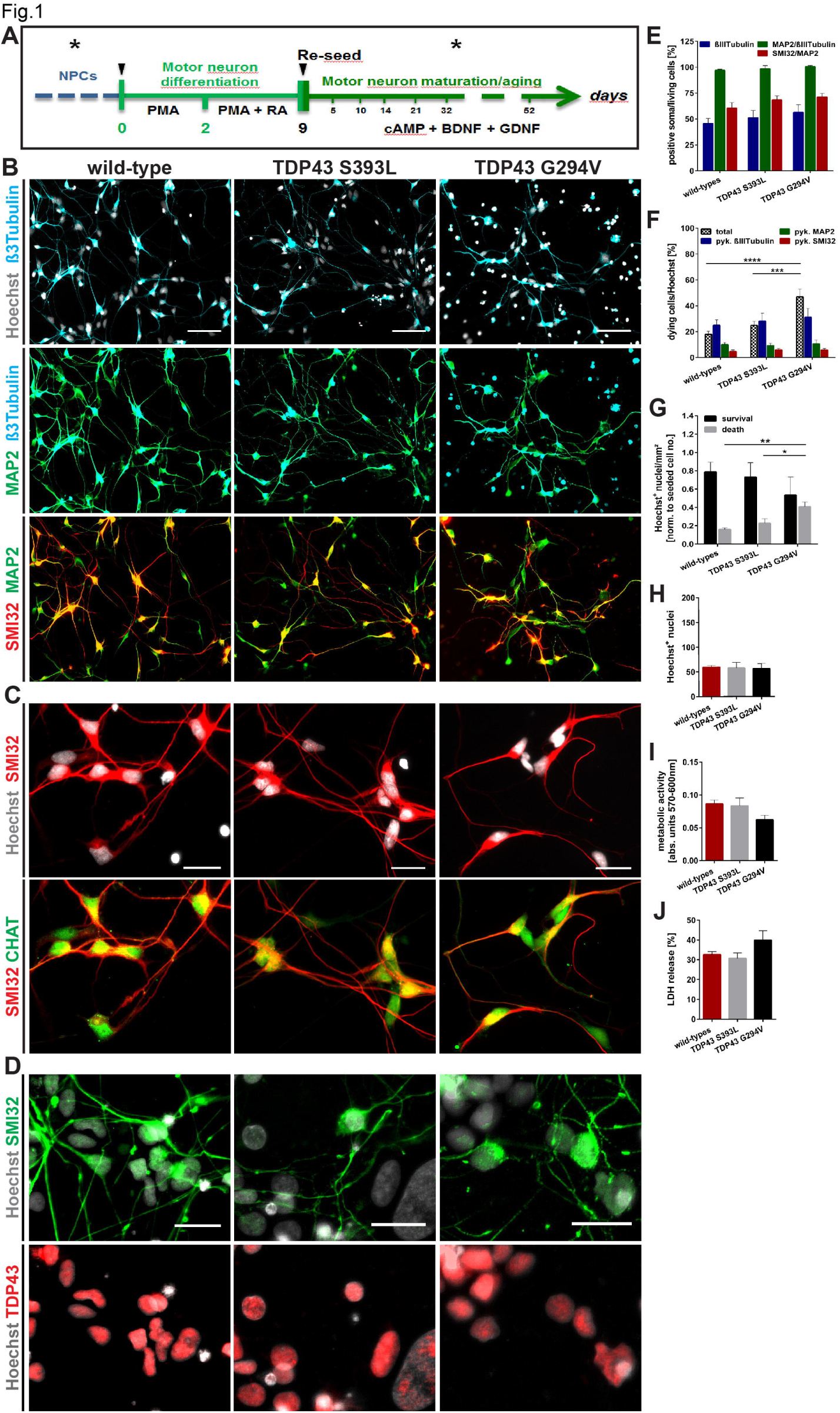
Characterization of spinal motor neuron differentiation potential. **(A)** Human spinal motor neuron differentiation steps. Black arrowheads indicate splits. Analysis (*) was performed at NPC stage and at several stages after re-seeding of the motor neuron cultures for maturation. **(B)** Representative immunofluorescence stainings of neurons generated from wildtype and TDP43 mutant patient NPCs 5 days post maturation induction. Shown are the pan-neuronal marker βIII-Tubulin, the mature neuron marker MAP2 and the spinal motor neuron marker SMI32. There was no difference in neuron morphology between wildtype and TDP43 mutant neuron types. Cell nuclei were stained with Hoechst. Scale bars: 75 µm. **(C)** Representative immunofluorescence stainings of mature motor neurons (SMI32) positive for the cholinergic spinal motor neuron marker choline acetyltransferase (CHAT). Cell nuclei were stained with Hoechst. Scale bars: 25 µm. **(D)** SMI32+ motor neurons showed no cytoplasmic accumulation of TDP43 protein. Scale bars: 25 µm. **(E)** Quantitative analysis of total βIII-Tubulin, MAP2/βIII-Tubulin and SMI32/MAP2 positive neuron soma as percentage of total living cell number (intact Hoechst (n=4). **(F)** Quantitative analysis of total cell death, βIII-Tubulin^+^ neuron death, MAP2^+^ neuron death and SMI32^+^ motor neuron death as percentage of total dying cell number (n=4); pyk. – pyknotic. **(G)** Quantitative analysis of the survival-and death rates of cells in wildtype and TDP43 mutant motor neuron cultures revealed increased death rates in TDP43 G294V mutant cultures when normalized to the initially seeded cell number (n=4). **(H)** Quantitative data of Hoechst^+^ nuclei count per image showed no differences in cell numbers between wildtypes and TDP43 mutant cultures (n=4-7). **(I)** Metabolic activity measurement by PrestoBlue^®^ assay resulted in similar behavior comparing wildtype and TDP43 mutant neuron culture metabolism (n=4-6). **(J)** Determination of cell damage by LDH activity measurement revealed no discrepancies between the wildtype and TDP43 motor neuron cultures (n=5-8). **(E-J)** Bar graphs show mean with SEM (1-way ANOVA (G-J) and 2-way ANOVA (E, F) with Bonferroni post hoc test, \**P*≤0.05, \*\**P*≤0.01, \*\*\**P*≤0.001, \*\*\*\**P*≤0.0001).

Total cell death and the number of dying neurons (marked by pyknotic cell nuclei, Figure 1 G,H) revealed a significantly increased total cell death in TDP43 G294V mutant motor neuron cultures (46.8±6.1 %) when compared to wildtype (17.9±2.4 %, *P*<0.0001) and TDP43 S393L mutant (24.8±3.1 %, *P*=0.0007). The death of young-(βIII-Tubulin), mature- (MAP2) or spinal motor neurons (SMI32) was not differing when wildtype and TDP43 mutants were compared.

Next we determined growth behavior and metabolism: Counting of Hoechst^+^ nuclei as well as metabolic activity and LDH release measurement showed no difference in growth of wildtype and TDP43 mutant motor neuron cultures (Figure 1 H-J, Hoechst: *P*=0.959, *F*=0.041; metabolism: *P*=0.126, *F*=2.513; LDH: *P*=0.130, *F*=2.341).

Together the early motor neuron maturation analysis showed an efficient homogeneous neuron differentiation with a high spinal motor neuron differentiation potential (SMI32^+^/CHAT^+^, Figure 1 C) in all analyzed motor neuron cultures.

Furthermore, both, wildtype and TDP43 mutant iPSC-derived motor neurons acquired normal neuronal function such as sodium and potassium currents (Figure 2 A-D), single and repetitive action potentials (Figure 2 E, F) as well as spontaneous postsynaptic currents (G-I) with no significant differences between healthy and TDP43 mutant motor neurons.

**Figure 2:**
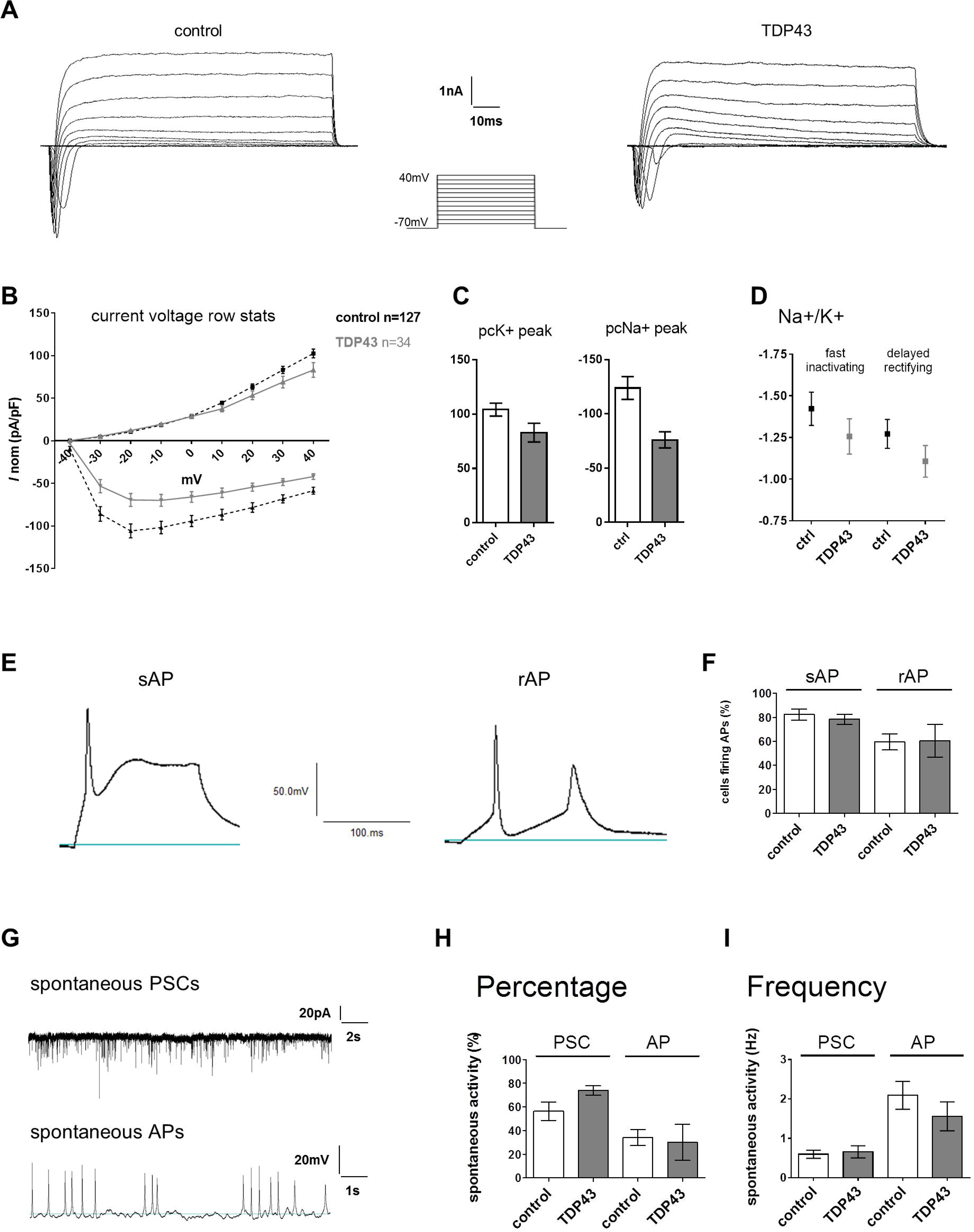
Functional maturation of healthy control and TDP43 mutant iPSC-derived motor neurons. **(A)** Stepwise depolarization in 10mV increments from a holding potential of −70mV to 40mV revealed voltage-dependent potassium and sodium currents **(B)** which were normalized for each respective cell capacitance. **(C)** No significant differences were observed in either potassium or sodium current peaks **(D)** nor sodium to potassium current ratios (Na^+^/K^+^) when comparing healthy control (n=127) to TDP43 mutant (n=34) iPSC derived neurons. **(E)** At least single (sAP) as well as repetitive trains of action potentials (rAP) were observed in the majority of the recorded neurons without a significant difference between groups **(F). (G)** Spontaneous activity was observed as post-synaptic currents (PSCs) and action potentials (APs). No differences between control and TDP43 mutant neurons were observed in the **(H)** percentage or **(I)** frequency of PSCs and APs.

### Mutant TDP43 motor neurons show age-related disturbed distal axonal organelle morphology

During aging mitochondria and lysosomes were visualized by Mito- and Lysotracker labelling of motor neuron cultures in directed microfluidic chambers (MFCs) at 2 and 4 weeks of maturation/aging (Figure 1 A & 3 A, B). Comparing wildtype and TDP43 mutants, measurement of neurite length variations from young to aged stages (i.e. fold-change over corresponding younger time point) revealed a constant proximal neurite mitochondrial network size for all lines whereas at the distal site wildtype cells exhibited a clear 1,179-fold increase (±0,194) as opposed to a moderate 0.721-fold decrease (±0,070) in the mutants (*P*=0.004) (Figure 3 D). Considering the number of measured organelle objects it could be excluded that the differences in neurite network size arose from different lysosome or mitochondria numbers (Figure 3 D; lysosomes: *P*=0.928; mitochondria: *P*=0.545). Moreover, there was no difference in the mitochondrial membrane potential between wildtypes and TDP43 mutant neurons (Figure 3 E).

**Figure 3:**
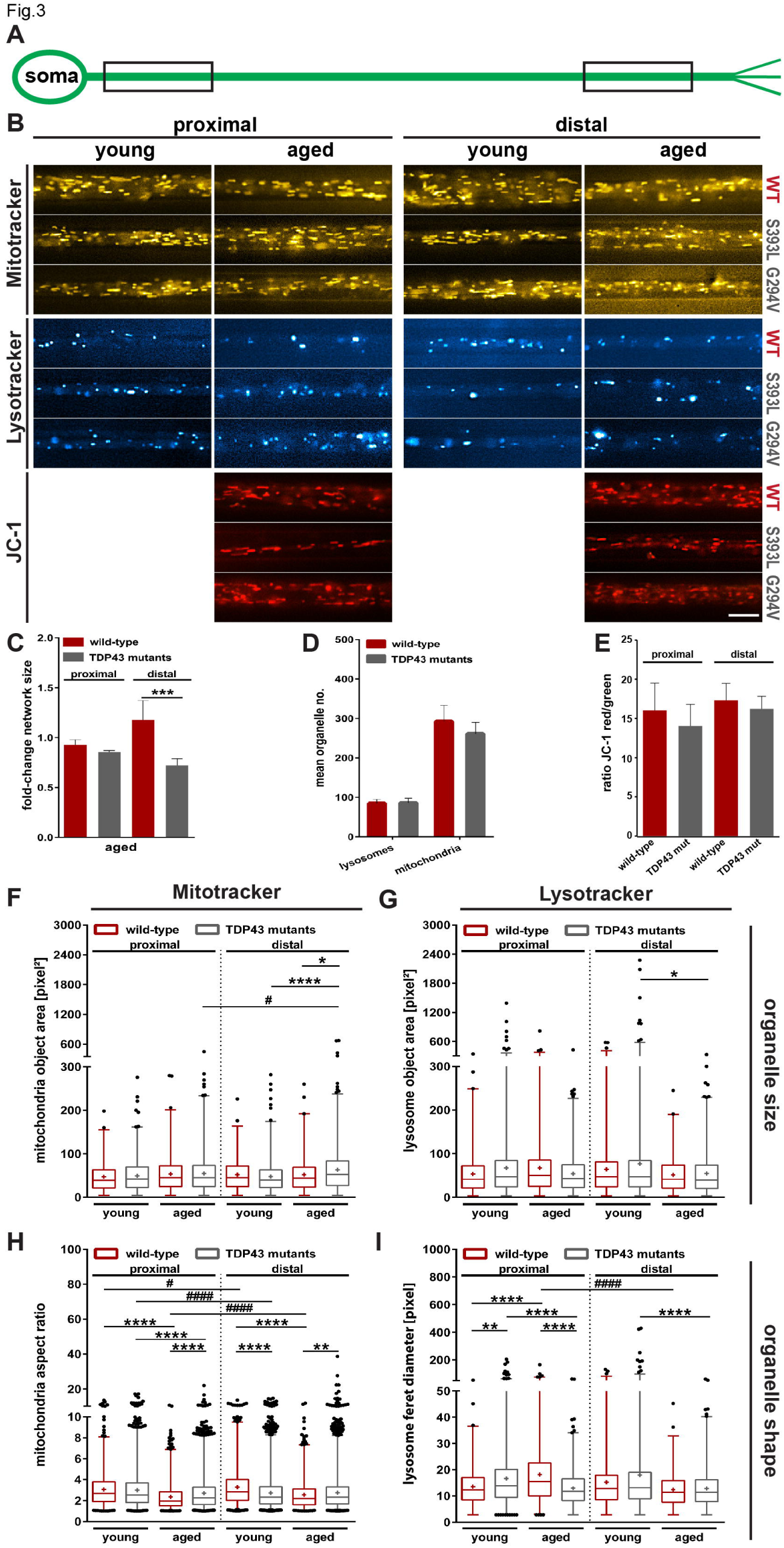
Cell organelle morphology and neurite network size during cellular aging. **(A, B)** Schematic illustration of analyzed proximal and distal sites of a motor neuron axon and representative images taken from the live cell movies of Mitotracker (upper panel, yellow), Lysotracker (middle panel, turquoise) and Mitotracker JC-1 (lower panel, red) in wildtype (WT), TDP43 S393L and G294V mutant microfluidic chamber (MFC) motor neuron cultures. Images show one passed MFC channel and display sections from 100x magnifications. Scale bars: 10 µm. **(C)** The proximal neurite network size (fold-change of aged cultures over corresponding young time point) did not differ between wildtype or TDP43 mutants. Conversely, the distal neurite network size decreased in the mutants over time as opposed to the increase in the wildtype. **(D)** The number of tracked organelles showed no difference between wildtype and TDP43 mutants. **(E)** No difference was seen in mitochondrial membrane potential. **(D-E)** Data of TDP43 mutants were pooled. Bar graphs show mean with SEM (n=3, 1-way ANOVA with Bonferroni *post hoc* test (D), Student’s t-test (E), \*\*\**P*≤0.001). **(F, G)** Quantification of organelle size of young and aged mitochondria (F) and lysosomes (G) revealed a severe distal mitochondrial phenotype in TDP43 mutant cultures characterized by increased size. **(H, I)** Quantification of organelle shape of young and aged mitochondria (H, aspect ratio) and lysosomes (I, Feret’s diameter) revealed significantly elongated mitochondria (increased aspect ratio) at both axon sites of aged TDP43 mutant neurons. **(F-I)** Data of TDP43 mutants were pooled. Box plots show mean (+) and median (line) and whiskers from 1-99 percentile (n=3-5, Kruskal-Wallis test with Dunn’s post hoc test; asterisks compare cell lines (e.g. wt/TDP43 mutants) within a condition (e.g. distal/proximal), rhombs compare between conditions within a cell line; *^/#^*P*≤0.05, **^/##^*P*≤0.01, \*\*\**P*≤0.001, ****^/#####^*P*≤0.0001). For detailed statistics see Suppl. Tables 2 - 5.

Mitochondria and lysosomes were both analyzed with respect to their individual organelle size and shape properties (Figure 3 F-I). The study of the mitochondrial area (=size) revealed a distal phenotype of aged TDP43 mutant MFC motor neurons whereas mitochondria at the proximal axon areas showed no differences. The analysis of lysosomal area resulted solely in a moderate distal phenotype of aged TDP43 mutant cultures when compared to young ones (Figure 3 G; detailed statistics are provided in Suppl. Tables 2 and 3). Together, the size of the majority of mitochondria and lysosomes were exclusively affected at distal axon sites of aged TDP43 mutant motor neuron cultures.

Next, we investigated mitochondrial and lysosomal shape (Figure 3 H, I). Our analysis of the mitochondrial aspect ratio (AR; indicates the shape ranging from round (AR=1) to elongated (AR>1) morphologies; long:short radius) revealed diverse phenotypes differing from proximal to distal axon sites (Figure 3 H). At proximal axons, we observed a mitochondrial shortening from young to aged cultures in both wildtype and TDP43 mutants that was also present in distal axons of wildtype neurons (proximal: 2.34±0.03, *P*<0.0001; distal: 2.54±0.03, *P*=0.003; Detailed statistics are presented in Suppl. Table 4). Taken together, the majority of mitochondria of TDP43 mutant MFC cultures revealed a global phenotype of imbalanced shape properties when proximal and distal axon sites were compared and in contrast to wildtype they got elongated during aging.

Furthermore, analyzing the Feret diameter of lysosomes (i.e. the diameter of the outer organelle’s perimeter) revealed a significant proximal decrease in TDP43 mutant motor neurons from young to aged stages as opposed to the increase in the wildtype during the same time. (Figure 3 I; detailed statistics are provided in Suppl. Table 5). Conversely, at the distal end we revealed a more vague trend towards smaller lysosome diameters in all lines.

### TDP43 mutant motor neurons show a dramatic loss of organelle motility during aging

The motility of mitochondria and lysosomes is a critical property especially in neurons since it mainly determines the energy supply and cellular waste removal of cells. We analyzed the organelle mean speed and the extent of organelle displacement in MFC motor neuron cultures and identified a drastic global (i.e. at either proximal and distal site) axonal transport deficiency in TDP43 mutants in aging but not young motor neurons (Figure 4 A; Suppl. Videos 1 & 2). In detail, mitochondria of TDP43 mutant neurons exhibited a drastic slow-down of mitochondrial mean speed at proximal and distal axons during aging (Figure 4 A, B; Suppl. Videos 1, 2; Suppl. Table 6). Consistently, we revealed a decrease of mitochondrial displacement (=extent of movement radius, measure for processivity in motility) (Figure 4 D, for detailed statistics see Suppl. Table 6; 8).

**Figure 4:**
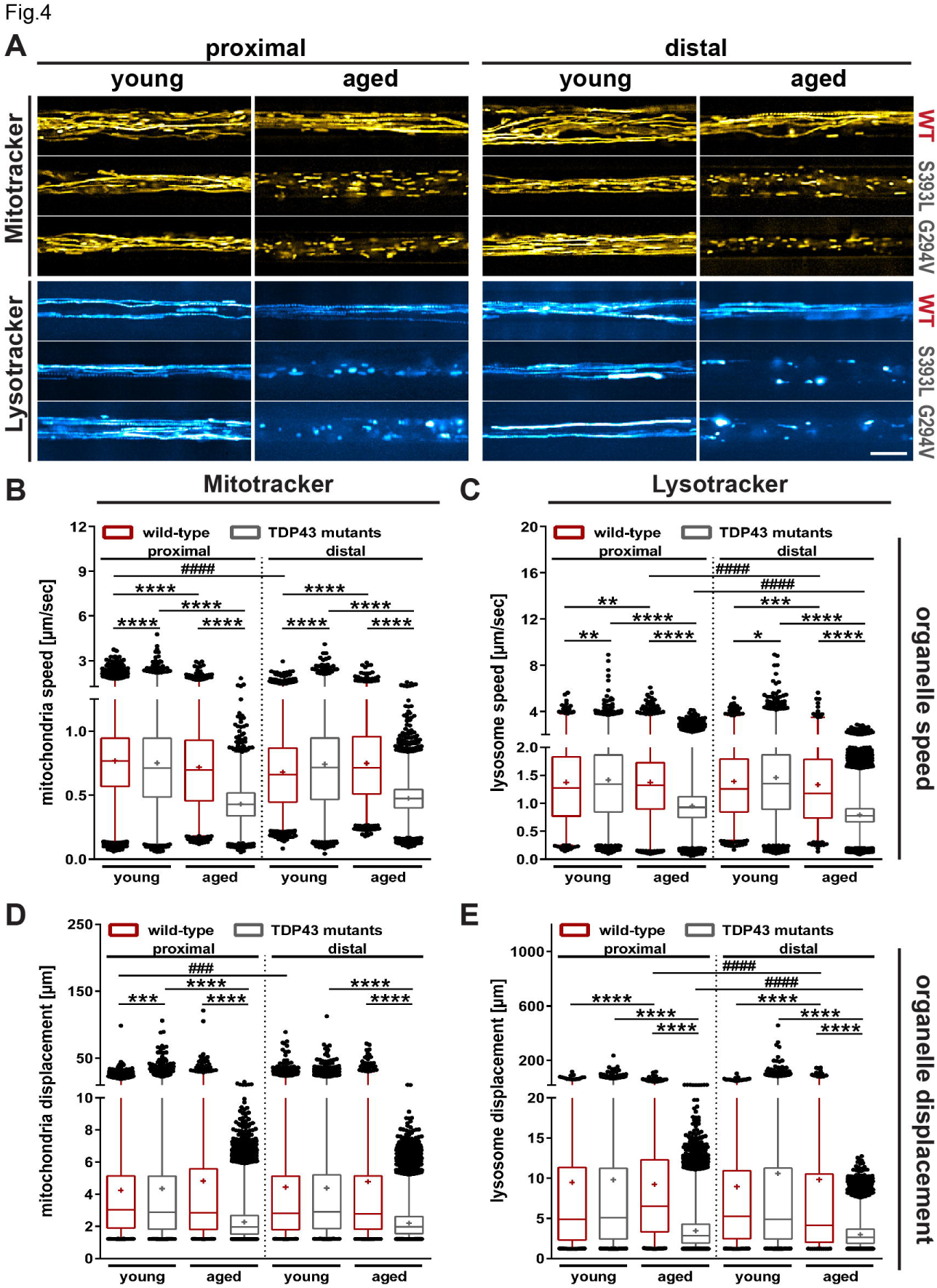
Cell organelle motility during cellular aging. **(A)** Shown are maximum intensity projections of entire movie stacks. Processively moving organelles (Mito-/Lysotracker) appear as continuous trajectories (bright lines) whereas less motile organelles as shorter lines and dots. **(B, C)** Organelle speed analysis of young and aged mitochondria (B) and lysosomes (C) compared in proximal and distal axon parts revealed a dramatic speed loss of either organelle type and axon sites in TDP43 mutant neurons. **(D, E)** Displacement analysis of young and aged mitochondria (D) and lysosomes (E) at proximal and distal axon sites revealed a dramatic decrease in processive motility of either organelle types and axon sites in aging TDP43 mutant motor neurons. **(B-E)** Data of TDP43 mutants were pooled. Box plots show mean (+) and median (line) and whiskers from 1-99 percentile (n=3-5, Kruskal-Wallis test with Dunn’s post hoc test; asterisks compare cell lines (e.g. wt/TDP43 mutants) within a condition (e.g. distal/proximal), rhombs compare between conditions within a cell line; \**P*≤0.05, \*\**P*≤0.01, \*\*\**P*≤0.01, ****^/####^*P*≤0.0001). For detailed statistics see Suppl. Tables 6-9.

The analysis of lysosomal mean speed uncovered similar trafficking defects (Suppl. Videos 1; 2): lysosomes of TDP43 mutant neurons significantly decreased their mean speed (Figure 4 A, C) and displacement (Figure 4 E, for detailed statistics see Suppl. Table 7, 9) at both axon sites during aging. Overall, our data indicate a gross decrease of mitochondrial and lysosomal organelle motility during aging at both distal and proximal axons in TDP43 mutant motor neurons.

### Aged TDP43 mutant motor neuron cultures show increased neuron death without signs of TDP43 protein aggregation formation

The neuronal homogeneity during the early neuron maturation stage (Figure 1) provided an ideal basis to study the cellular development of aging wildtype and TDP43 mutant motor neurons at later time points as analyzed by live cell imaging. In order to investigate if the used human model systems in this study recapitulate ALS hallmarks of neuron death and degeneration the survival of neurons including spinal motor neurons was investigated at different time points during cellular aging. The analysis from young (day 5 of maturation) to aged (day 32 of maturation) mutant cultures showed a significant loss in the total amount of βIII-Tubulin (Figure 5 A; day 5: 53.69 ± 4.83 %, day 32: 19.53 ± 3.14 %; *P*<0.0001), MAP2 (Figure 5 B; day 5: 53.49 ± 5.95 %, day 32: 33.40 ± 5.93 %; *P*=0.022) and SMI32 positive neurons (Figure 5 C; day 5: 34.67 ± 4.10 %, day 32: 5.0 ± 1.05 %; *P*<0.0001) whereas in the wildtype this loss was only moderate (Figure 5 A-C; day 32: βIII-Tubulin: 52.51 ± 6.94 %, *P*=0.0002; MAP2: 56.03 ± 3.62 %, *P*=0.013; SMI32: 15.93 ± 2.50 %, *P*=0.004). Together, our results document a clear vulnerability and compromised survival in TDP43 mutant spinal motor neurons during cellular aging whereas wildtype cells could maintain a constant neuron pool. Interestingly, the age-dependent increase of neuronal death was not accompanied by TDP43 protein aggregation (Figure 1 D). Wildtype as well as TDP43 mutant cells showed a dominant nuclear localization of TDP43 protein and no detectable phosphorylated TDP43(Supplemental Figure 1).

**Figure 5:**
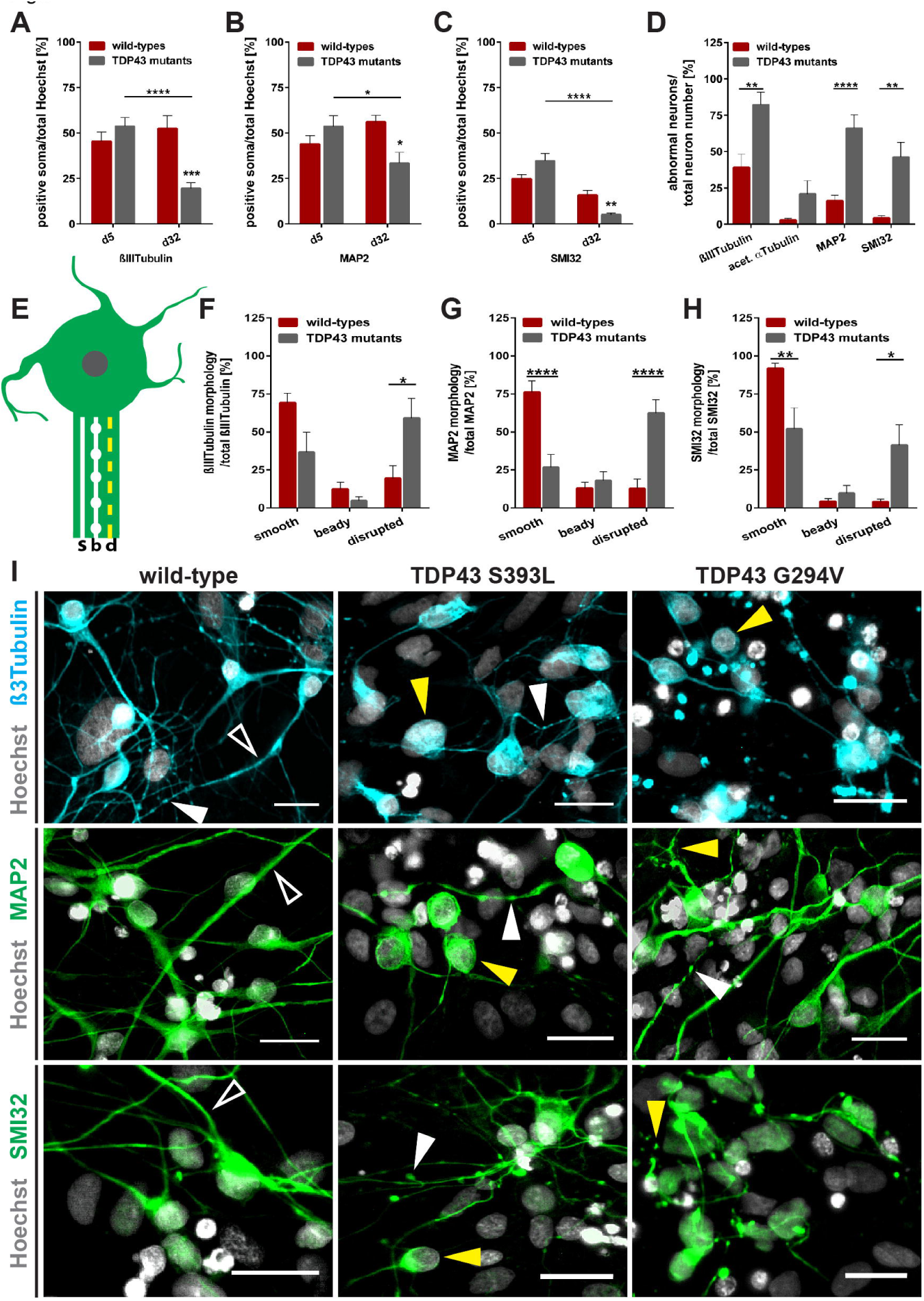
The TDP43 mutation related loss of neurons during aging is accompanied by abnormal neurofilament morphologies. **(A-C)** The total number of βIII-Tubulin (A), MAP2 (B) and SMI32 (C) positive neurons was counted in young (day 5) and aged (day 32) maturation cultures and revealed a severe loss of every type of neuron in aged TDP43 mutant cultures, (n=6-8). **(D)** The number of abnormal neurons was evaluated in aged cultures and revealed a highly increased number of misshaped βIII-Tubulin, MAP2 and SMI32 positive neurons indicated by beady (b) and disrupted (d) cytoskeleton morphologies. **(E)** Schematic demonstration of a neuron forming smooth (s), beady (b) and disrupted (d) cytoskeleton morphologies, (n=3-8). **(F-H)** The number of neurons showing smooth, beady or disrupted cytoskeleton per neuron type was calculated and revealed a dramatic loss of normal smooth cytoskeleton morphology for MAP2 (G) and SMI32 (H) positive mature neurons in TDP43 mutants, consistent with a drastic increase of disrupted cytoskeleton morphology in all neuron types in TDP43 mutants (n=6-8). **(I)** Representative immunofluorescence images of cytoskeletal morphologies (smooth: hollow white arrow heads, beady: filled white arrow heads, disrupted: filled yellow arrow heads) of 32 days maturated wildtype and TDP43 mutant motor neurons. Cell nuclei were stained with Hoechst. Images display sections of 20x magnifications. Scale bars: 25 µm. **(A-D and F-H)** Data of wildtype 1-3 as well as TDP43 mutant lines were pooled. Bar graphs show mean with SEM (2-way ANOVA with Bonferroni *post hoc* test, *P≤0.05, **P≤0.01, ***P≤0.001, ****P≤0.0001).

### TDP43 mutant neurons show dramatic axo-skeletal degeneration during aging

The analysis of cytoskeletal alterations during neuronal aging revealed dramatic differences between wildtype and TDP43 mutant motor neuron cultures (Figure 5 D-I). We analyzed tubulin (βIII-Tubulin), microtubule-associated protein Tau (MAP) as well as neurofilament-H (NF-H, SMI32). Cytoskeletal proteins of wildtype neurons appeared as firm, smooth and thick (indicated as “smooth”). In contrast, the cytoskeleton of TDP43 mutant neurons exhibited disrupted morphologies of shortened and fragile dendrites (MAP2) and axons (SMI32) combined with striking loss of neurites and branching (indicated as “disrupted”). A third structural morphology change identified as smooth filament with intermittently occurring filament beads (indicated as “beady”) could be detected in wildtype cells and TDP43 mutants. A schematic demonstration is given in Figure 5 E and representative images are shown in Figure 5 I. Morphologically aberrant neurons were quantified at day 32 of maturation (Figure 5 E-H) and the cytoskeleton structures were classified by their appearance (smooth, beady and disrupted; Figure 5 E). Collectively, our findings document an age- and TDP43-mutation-related abnormal cytoskeletal morphology in mature spinal motor neurons.

### D-Sorbitol rescues mutant TDP43 motor neuron organelle morphology and motility without inducing TDP43 aggregation

Osmolytes are small organic compounds that affect protein stability and are ubiquitous in living systems. D-sorbitol is known as protein protecting and stabilizing osmolyte ^31^. Furthermore, D-sorbitol was reported to recruit TDP43 to stress granules ^18^, to induce cytoplasmic mislocalization and to interfere with the quality control machinery in TDP43 mutants ^19^. To further address these findings, we treated aged MFC motor neuron cultures with 40mM up to 400mM of D-sorbitol for 24h. Interestingly, D-sorbitol did not induce any cytoplasmic TDP43 protein localization or aggregation (Figure 6 K, L; Supplement Figure 2).

**Figure 6:**
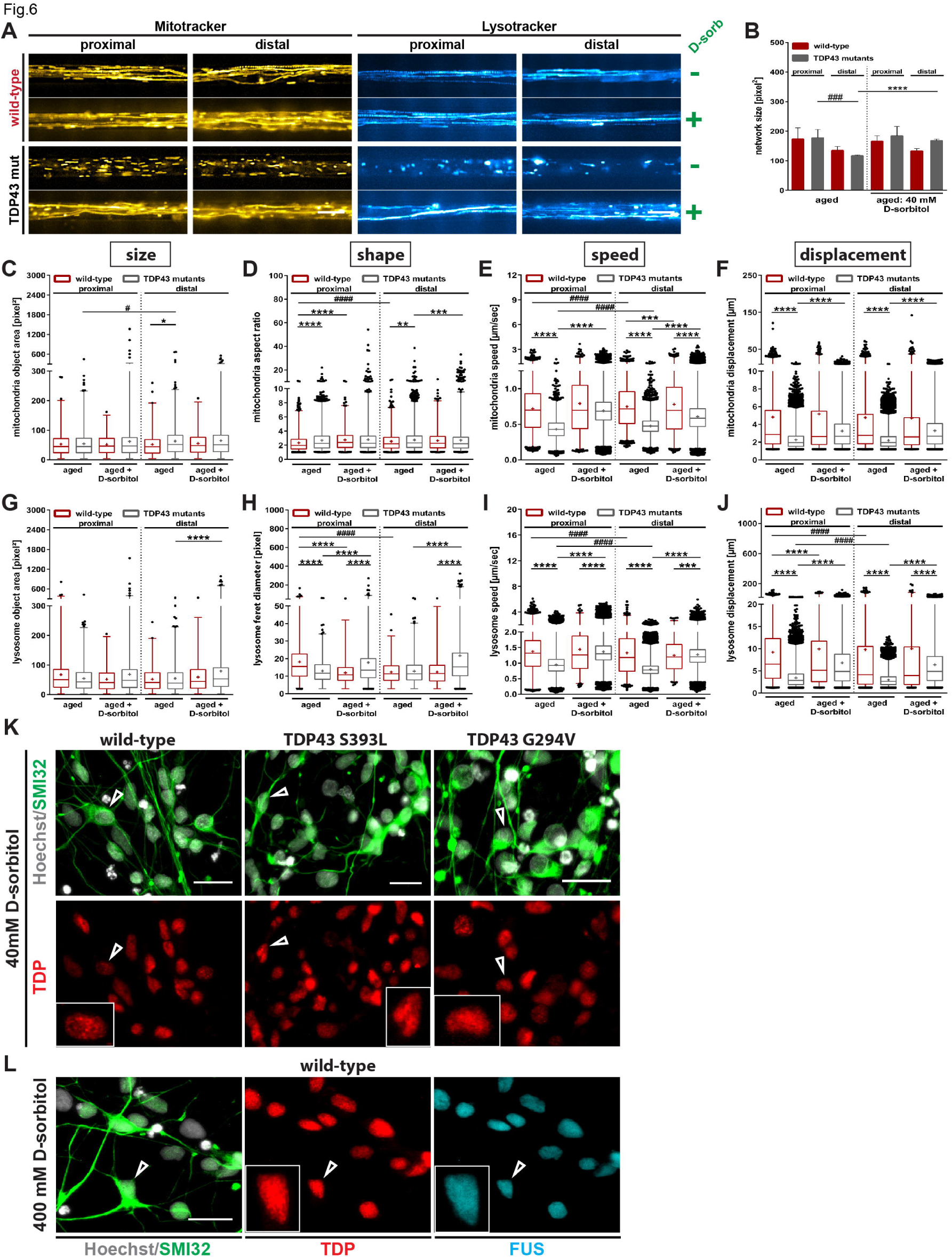
Analysis of cell organelles in aged spinal motor neuron cultures after 24 h D-sorbitol treatment. **(A)** Representative images taken from live cell movies of Mitotracker (left) and Lysotracker (right)^38^ in D-sorbitol-treated wildtype (WT), TDP43 S393L and G294V mutant MFC motor neuron cultures. Images show one passed MFC channel at proximal and distal sites and display sections from 100x magnifications. Scale bars: 10 µm. **(B)** Reduced neurite network size in aged TDP43 mutants at the distal axon site was restored by the treatment with 40mM D-sorbitol. **(C, G)** Proximal and distal organelle sizes of mitochondria (C) and lysosomes (G) from aged and D-sorbitol treated MFC motor neuron cultures showed compensation of the observed distal differences of mitochondrial size between wildtype and TDP43 mutants and displayed a lysosomal size increase in TDP43 mutants. **(D, H)** Proximal and distal shape analysis of aged mitochondria (D) under D-sorbitol treatment showed a mimic of the proximal mitochondrial elongation of TDP43 mutants in the wildtype. For lysosomes (H) the proximal phenotype got reversed and the TDP43 mutants were strongly influenced at the distal axon site. **(E, I)** Proximal and distal motility of mitochondria (E) and lysosomes (I) was rescued to normal speed in TDP43 mutants close to wildtype levels at both axon sites through D-sorbitol. **(F, J)** Dito for organelle displacement. **(K, L)** Representative immunofluorescence images of D-sorbitol-treated aging motor neurons negative for cytoplasmic TDP43 aggregation. **(K)** Wildtypes and TDP43 mutants showed no cytoplasmic TDP43 protein aggregation after treatment with 40 mM D-sorbitol for 24h. **(L)** Increasing the D-sorbitol concentration to up to 400 mM did not induce cytoplasmic TDP43 or FUS mislocalization/aggregation in wildtype motor neurons. Cell nuclei were stained with Hoechst. White arrowheads mark a single representative motor neuron. Scale bars: 25 µm. **(B-J)** Data of TDP43 mutants were pooled. Bar graphs (B) show mean with SEM. Box plots (C-J) show mean (+), median (line) and whiskers from 1-99 percentile (n=3-5, Kruskal-Wallis test with Dunn’s *post hoc* test; asterisks compare cell lines (e.g. wt/TDP43 mutants) within a condition (e.g. distal), rhombs compare between conditions within a cell line; *^/#^*P*≤0.05, \*\**P*≤0.01, ***^/###^*P*≤0.001, ****^/####^*P*≤0.0001). For detailed statistics see Suppl. Tables 2-9.

Very surprisingly, however, we found a profound rescue of organelle morphology and deficient motility in mutant TDP43 motor neurons by D-sorbitol treatment (40 mM). Specifically, D-sorbitol caused a significant increase of the distal axon TDP43 mutant network size (Figure 6 A, B; untreated: 117.2±2.93 pixel², D-sorbitol: 168.0±5.62 pixel²; *P*<0.0001). Considering the single organelle size the treatment with D-sorbitol eliminated the soft differences in distal mitochondria size between aged wildtype and TDP43 mutant MFC cultures (Figure 6 C; treated wildtype: 55.3 ± 2.89 pixel², treated TDP43 mutants: 65.0 ± 3.35 pixel²; *P*>0.999). For lysosomes it led to a restoration of the young distal TDP43 mutant morphology (see Figure 3 G) by highly increased lysosomal organelle area mean to 79.24 ± 4.08 pixel² (Figure 6 G; untreated: 54.57 ± 1.83 pixel², P<0.0001) whereas the wildtype stayed unaffected. Together D-sorbitol affected organelle sizes only at distal axon sites whereas the organelle shape and motility were influenced at both axon sites.

Additionally, D-sorbitol showed a clear rescue of organelle motility (Figure 6 E, F, I and J and Suppl. Videos 3 & 4), i.e. mitochondrial speed (Figure 6 E) and displacement (Figure 6 F), as well as lysosomal speed (Figure 6 I) and displacement (Figure 6 J; for detailed statistics see Suppl. Tables 2 - 9).

## Discussion

Here we show age-dependent neurodegeneration of patient-derived TDP43 mutant motor neurons similar to Bilican and colleagues ^12^. Furthermore, we present substantial data arguing against TDP43 aggregation as upstream event in the pathophysiology of TDP43-ALS. Initially, morphologically healthy appearing motor neurons (Figure 1, similarly reported in ^11^) revealed robust TDP43-mutation-dependent abnormal phenotypes of mitochondria and lysosomes (Figures 3, 4, 6) and the cytoskeleton (Figure 5) during aging without signs of cytoplasmic TDP43 aggregation (Figures 1, 5).

Since the first description of a motor neuron disease a well-known hallmark in the pathology is progressive motor neuron degeneration in the spinal cord ^32^ and, by modern neuropathology techniques, pathogenic TDP43 aggregation in motor neurons of ALS and FTLD ^5,33^ and other neurodegenerative disorders ^34^. In this study, we document pathological degeneration of *in vitro* cultivated patient-derived (iPSCs) spinal motor neurons of TDP43 mutation carriers characterized by dramatic neuron and spinal motor neuron loss during aging and the formation of severely abnormal cytoskeletal structures in mutant cultures (Figure 5). In contrast, we could not observe cytoplasmic TDP43 localization or aggregation in aged neurons and spinal motor neurons (Figure 1, 5) assuming a TDP43 protein mislocalization-independent mechanism of neurodegeneration. We cannot rule out that the mutations used in the current study behave slightly different to other TDP43 mutations ^35^. However, our data is not in contrast to recent studies reporting cytosolic mislocalization of TDP43 since, on a closer look, also in these models suing different TDP43 mutations only up to one third of cells showed cytosolic mislocalization while the remaining cells still showed physiological nuclear TDP43 ^10,11,14^.

Surprisingly, we could not detect relevant cytoplasmic TDP43 mislocalization or aggregation even after treatment with the osmolyte D-sorbitol being reported to induce stress granules formation and cytoplasmic mislocalization of TDP43 ^18^. Furthermore, D-sorbitol rescued organelle morphology and motility in TDP43 mutant patient-derived neurons (Figure 5). Neurite outgrowth was reported being reduced in murine models of TDP43 ^9,20^ which could be restored by non-toxic proteasome inhibition ^20^. D-sorbitol is known as protein protecting and stabilizing osmolyte ^31^. This could explain our rather unexpected results showing that the osmolyte sorbitol leads to restoration of the axon trafficking phenotype. Together, both results argue against a TDP43 aggregation-dependent mechanism as early event in neurodegeneration of TDP43 motor neuron degeneration.

The size of the majority of mitochondria and lysosomes was mainly affected at distal axon parts (directed to axon endings) indicated by elongated mitochondria and shrunken lysosomes in comparison to wildtype cases (Figure 3 & 6). Furthermore, we detected in aged but not young TDP43 mutant motor neurons an overall organelle trafficking deficit (Figure 4 & 6). This is in accordance with TDP43 mice model data showing lack of mitochondria in motor axon terminals ^10^. Furthermore, TDP43 was reported to impair mitochondria and lysosome morphology and function in cell models of ALS. Interestingly, both overexpression and knockdown caused a phenotype in primary murine neurons suggesting a tight regulation of TDP43 ^22,23^.

TDP43 was shown to be actively transported in the axon thereby forming cytoplasmic mRNP granules that undergo bidirectional, microtubule-dependent transport in neurons and facilitate delivery of target mRNA to distal neuronal compartments ^14,15^. TDP43 mutations impair this mRNA transport function ^14,15^. Several lines of evidence suggest that mRNA encoding cytoskeletal proteins is cargo of such mRNA-transporting proteins being locally translated in the distal axon ^36,37^. For instance, neurofilament light chain (NEFL) mRNA was shown to be transported in such TDP43 granules and was reduced in TDP43 mutant motor neurons ^10,15^. These data fit to our observations of severely disturbed cytoskeletal proteins such as tubulin, microtubules and neurofilaments. Furthermore, this could induce a vicious cycle by additionally impairing microtubule-dependent mRNP granule transport ^14,15^.

Together, we here report typical age-related motor neurodegeneration in TDP43 mutant ALS patient-derived motor neurons leading to a severe organelle transport deficiency, cytoskeletal disturbance and neurodegeneration without TDP43 protein mislocalization or aggregation formation. This challenges the current concept of protein aggregation as the main cause of motor neuron disease pathology and calls for a search for therapeutic targets independent of TDP43 aggregation.

### Conclusions

We here show aging-related neurodegeneration in patient specific TDP43 mutant iPSC-derived motor neuron cultures. Cell organelles as mitochondria and lysosomes showed TDP43 mutation dependent abnormalities in size, shape, speed and motility which all appeared without TDP43 mislocalization or aggregation formation. Finally, D-sorbitol – known to induce stress granules and cytoplasmic mislocalization of TDP43 – rescued axonal trafficking phenotypes without any signs of TDP43 mislocalization or aggregation formation. Our data suggest TDP43 mutation – dependent but cytosolic aggregation-independent mechanisms of motor neuron degeneration in TDP43 ALS.

## Additional information Authors’ contributions

Nicole Kreiter, geb. Wächter: Conception and design, collection and assembly of data, data analysis and interpretation, manuscript drafting.

Arun Pal: Data collection and assembly, partial analysis and interpretation, manuscript drafting.

Xenia Lojewski: Generation of iPS and derivation of NPCs from patient fibroblasts, manuscript drafting.

Philippe Corcia: TDP43 mutant patient fibroblast delivery, manuscript drafting.

Maximilian Naujock: Data collection and assembly, partial analysis and interpretation, manuscript drafting.

Peter Reinhardt: Generation of iPS and derivation of NPCs from patient fibroblasts, manuscript drafting.

Jared Sterneckert: Generation of iPS and derivation of NPCs from patient fibroblasts, manuscript drafting.

Susanne Petri: Data collection and assembly, partial analysis and interpretation, manuscript drafting.

Florian Wegner: Data collection and assembly, partial analysis and interpretation, manuscript drafting.

Alexander Storch: Conception and design, data interpretation, manuscript drafting.

Andreas Hermann: Conception and design, principal investigator, collection and assembly of data, data interpretation, manuscript drafting and final approval.

## Acknowledgements

We thank the patients and controls for skin biopsy donation. We acknowledge the help in cell culture by Sylvia Kanzler and Katja Zoschke. The work was supported in part by the Helmholtz Virtual Institute (VH-VI-510) “RNA dysmetabolism in ALS and FTD” for the funding of the project to A.H. and A.S., the “Deutsche Gesellschaft für Muskelerkrankungen (He 2/2)”, the MeDDrive program of the Medical Faculty at the Technische Universität Dresden to A.H., the Center for Regenerative Therapies Dresden, the Roland Ernst Stiftung Saxony to A.H., BIOCREA GMBH to A.H. and the NOMIS foundation to A.H..

